# Revised testing procedures do not elicit magnetoreceptive behavior in *C. elegans*

**DOI:** 10.1101/349944

**Authors:** Lukas Landler, Gregory C. Nordmann, Simon Nimpf, Tobias Hochstoeger, Daniel Kagerbauer, David A. Keays

## Abstract

A diverse range of species is known to rely on the Earth’s magnetic field for spatial information. Vidal-Gadea et al. claimed that *C. elegans* are magneto-sensitive, exploiting the magnetic field to guide their burrowing behavior [1]. Our attempts to replicate their findings were unsuccessful [2], which Vidal-Gadea attributed to the satiety of the animals and the environment in which they were raised. Here, we report our repeated experiments, having adopted several suggestions proposed by Vidal-Gadea et al. [3]. We find that shortening the length of the behavioral assay and raising the animals in a Faraday cage does not result in magnetotactic behavior. We reluctantly conclude that the assays employed by Vidal-Gadea are not robust or *C. elegans* are not magneto-sensitive.

## Introduction

It has recently been reported by Vidal-Gadea and colleagues that *Caenorhabditis elegans* are magneto-sensitive [1]. In making this assertion they employed three assays: (1) a vertical burrowing assay; (2) horizontal plate assay; and (3) a magnetotaxis assay. In the latter assay, ~50 worms are placed on the center of a petri dish and are allowed to choose between two goal areas, one with a magnet underneath and one with a control disc. Worms reaching the goal areas are immobilized by the sodium azide, counted, and a preference index is calculated. While Vidal-Gadea et al. [1] reported that worms were significantly attracted to the magnet (e.g. n = 36 plates, t-test, p < 0.001), we observed no differences in the distribution of worms (n = 49 plates, Wilcoxon signed rank test, V = 565, n. s). In response to our experiments Vidal-Gadea et al. [3] raised four main concerns. First, we adapted the time of the magneto-taxis assay (increasing it from 30 min to 1h) enabling more worms to reach the goal area. They argued that this modification could lead to a shift in the satiation state of the worms (from fed to starved), altering their magneto-sensitivity. Second, we raised our worms in a standard incubator which generates electromagnetic noise. Vidal-Gadea and colleagues proposed that this might alter the magnetic response. Third, they suggested that our magneto-taxis results should have been compared to a control experiment, where control brass coins are placed under both goal areas. Finally, they argued that worms should be freshly thawed for the experiments as they might loose magnetoreceptive abilities after many generations in a lab environment. In this manuscript we address these issues, once again performing our experiments in our mu metal-shielded room employing strict blinded quantitation. We report that modification of the aforementioned parameters do not result in magnetotactic behavior in worms.

## Results

### Magneto-taxis assay

In light of the response of Vidal-Gadea et al. [3] we replicated our experiments adopting the measures they suggested. We found that reducing the time of the magneto-taxis assay from 1h to 30 min, did not induce magnetotactic behavior. The preference index was not significantly different from zero (Figure 1A, Wilcoxon signed rank test, n = 20 plates, V = 98.5, n. s.), nor were these results different from control trials that employed two brass discs without a magnet (Figure 1B, Mann-Whitney U-test, n = 20 plates, U = 182, n. s.).

Next we assessed whether raising worms in an incubator might be problematic. We therefore grew worms in a faraday cage (to minimize electromagnetic noise) at a fixed temperature (20°C). We performed the magnetotactic assay (for 30mins) and calculated the preference index. Worms exhibited no preference for the goal area associated with the magnet (Figure 1C, Wilcoxon signed rank test, n = 20 plates, V = 91, n. s.)). We compared this experimental group to a no-magnet control, and again did not observe a significant difference between groups (Figure 1D, Mann-Whitney U-test, n = 20, U = 190.5, n. s.).

**Figure 1:**
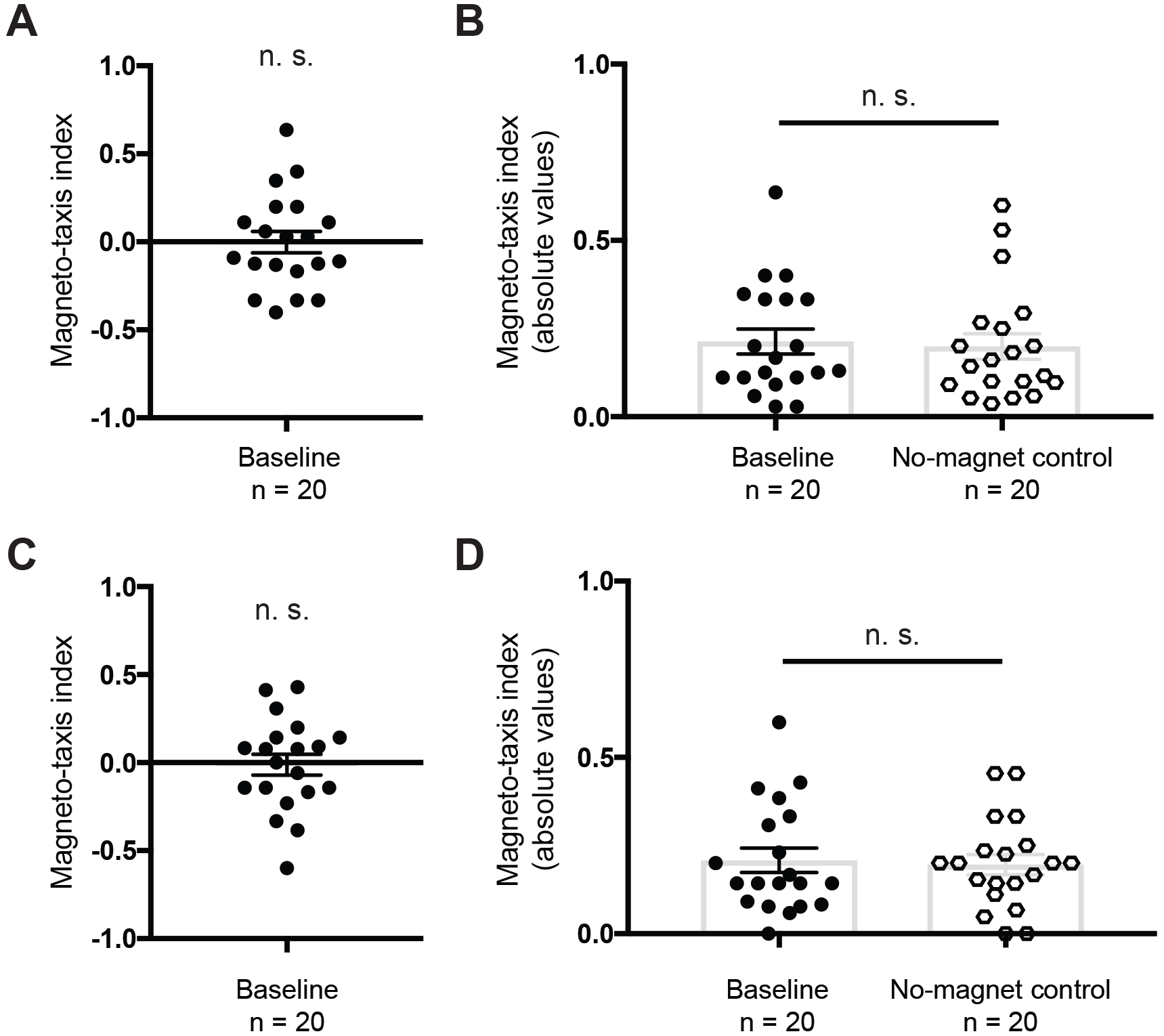
Results from the magneto-taxis experiment. **(A)** Worms were tested on a plate with a magnet and a brass control underneath two different goal areas (baseline) for 30 min, and the magneto-taxis index calculated. Animals did not show a preference for the magnet (n = 20 plates, Wilcoxon signed rank test, V = 98.5, n. s.). **(B)** Brass control discs were placed under both goal areas, and the distribution of worms compared to the results obtained for **(A)**. For this comparison the data has been plotted showing the absolute values of the magneto-taxis index (i.e. all values are positive). We did not observe a significant difference when comparing these two groups (Mann-Whitney U-test, n = 20, U = 182, n. s.). **(C)** Worms raised in a Faraday cage to reduce exposure to electromagnetic noise did not show a preference for the goal area above the magnet when tested for 30 min (Wilcoxon signed rank test, V = 91, n. s.). **(D)** Similarly, the baseline experiment did not differ significantly from the no-magnet control, when plotting the absolute values (Mann-Whitney U-test, n = 20, U = 190.5, n. s.).

## Discussion

We adopted the suggestions proposed by Vidal-Gadea et al. [3], and changed the trial time, the environment that worms are raised, and used freshly thawed worms. None of these changes resulted in a significant preference for the magnet over a brass control. Moreover, the distribution of worms was similar to the negative control experiments with brass coins only. Our results demonstrate that neither the satiety nor electromagnetic noise in the standard worm incubators explain the lack of a magnetic response. It should be further noted that other investigators have been unable to replicate the findings of Vidal-Gadea (Robert Fitak, personal communication). We reluctantly conclude that the assays devised by Vidal-Gadea et al. are not suitable as a robust method to decipher the physical mechanism and molecular machinery of magnetoreception in animals.

## Acknowledgement

Our thanks go to: Boehringer Ingelheim for funding basic research at the Research Institute of Molecular Pathology; the IMP graphics department and the media kitchen; Manuel Zimmer, Annika Nichols and Richard Latham for assistance with C. elegans work and E. Pascal Malkemper for help with the manuscript. The C. elegans strain was provided by the CGC which is funded by NIH Office of Research Infrastructure Programs (P40 OD 010440).

## Methods and material

### Animals

*C. elegans* (N2, obtained from Caenorhabditis Genetics Center) were kept on the *Escherichia coli* strain OP50. For the baseline magneto-taxis experiment animals were maintained in incubators in the dark at 20°C. For the ‘electromagnetic noise free’ magneto-taxis assay we placed the culture plates in a copper Faraday cage in a room that was kept on 20°C. We used never starved adult hermaphrodite worms for all assays. Prior to the experiments worms were synchronized (bleached). Worms were tested within 10 min after removal from the culture plate (satiation state ‘fed’).

### Magneto-taxis assay

We followed the experimental procedure described previously [2], however adding a few changes (reducing the trial time to 30 min, adding a no-magnet control trial and cultivating worms for the second experiment in a Faraday cage).

Worms (~ 50) were placed in the center of an agar (3%) filled 100mm style petri dish. Two goal areas (‘scoring’ circles) were marked in equidistance to the center of the petri dish and a magnet (N42 Neodymium 3.5 cm) was placed (north up) under one of the scoring circles and a brass control (same dimensions as the magnets) under the other one. The plates were then covered with aluminum foil and put in an electromagnetically shielded room (mu-metal and aluminum shielding against static and oscillating magnetic fields, see Landler et al. [2]). After 30 minutes the number of worms on either scoring circle were counted. In parallel we performed a control experiment in which we had brass control coins under both scoring circles. The experiments were performed in a double blind manner, i.e. the person counting the worms did not know under which scoring circle the magnet was, nor was he aware if the plate was a no-magnet control plate. For the plates with a magnet we calculated a magneto-taxis index (MI), which was the number of worms on the side with the magnet (M) subtracted by the worms on the control side (C) and divided by the total number of scoring worms (MI = (M−C)/(M+C)). For the control only trials we calculated a control index by randomly choosing one of the two sides as the reference and then taking the absolute value to compare it to the absolute values of the magneto-taxis index. In order to test if the preference index was higher than zero we used a one-tailed Wilcoxon one-sample test. We tested if the absolute values of the preference index and the control index differed using a one tailed Mann-Whitney U-test.

